# Slower environmental change hinders adaptation from standing genetic variation

**DOI:** 10.1101/203760

**Authors:** Thiago S. Guzella, Snigdhadip Dey, Ivo M. Chelo, Ania Pino-Querido, Veronica F. Pereira, Stephen R. Proulx, Henrique Teotónio

## Abstract

Evolutionary responses to environmental change depend on the time available for adaptation before environmental degradation leads to extinction. Explicit tests of this relationship are limited to microbes where adaptation depends on the order of mutation accumulation, excluding standing genetic variation which is key for most natural species. When adaptation is determined by the amount of heritable genotype-by-environment fitness variance then genetic drift and/or maintenance of similarly fit genotypes may deter adaptation to slower the environmental changes. To address this hypothesis, we perform experimental evolution with self-fertilizing populations of the nematode *Caenorhabditis elegans* and develop a new inference model that follows pre-existing genotypes to describe natural selection in changing environments. Under an abrupt change, we find that selection rapidly increases the frequency of genotypes with high fitness in the most extreme environment. In contrast, under slower environmental change selection favors those genotypes that are worse at the most extreme environment. We further demonstrate with a second set of evolution experiments that, as a consequence of slower environmental change, population bottlenecks and small population sizes lead to the loss of beneficial genotypes, while maintenance of polymorphism impedes their fixation in large populations. Taken together, these results indicate that standing variation for genotype-by-environment fitness interactions alters the pace and outcome of adaptation under environmental change.

## Introduction

With human activities predicted to increase rates of climate change (Stocker et al. 2013), it has become urgent to pinpoint the ecological and evolutionary conditions by which natural populations survive and adapt at different rates of environmental change. It is generally accepted that low rates of environmental change allow more time for new beneficial mutations to appear and, consequently, to promote adaptation and to rescue populations from extinction (Lynch and Lande 1993, Kopp and Hermisson 2009, Lande 2009, Chevin et al. 2010). Experimental evolution results from studies of with microbes support this idea (Perron et al. 2008, Collins and de Meaux 2009, Bell and Gonzalez 2011, Gorter et al. 2015), with one study in particular having found that population survival and adaptation depend on the order of mutation accumulation and epistasis for fitness (Lindsey et al. 2013). However, most species in nature have small populations, are genetically structured by geography or reproduction system, have long generation times and/or are unable to migrate to their favored habitats. In all these cases, survival and adaptation to changing environments will depend on pre-existing genetic diversity, and less so on mutation accumulation (Hill 1982, Matuszewski et al. 2015).

Adaptation to changing environments from pre-existing genetic diversity is conditional on how each genotype performs within the environments that may be encountered in the near future (Fig. 1A). Depending on the shape of these “fitness reaction norms” (Chevin et al. 2010, Walsh and Lynch 2014, Gorter et al. 2015), and evolutionary history (Lande 2009, Gonzalez and Bell 2013), natural selection may initially favor genotypes at intermediate stressing environments that are not necessarily the best at the more extreme environments. Short-term adaptation will therefore be determined by the amount of heritable genotype-by-environment fitness variance. Two predictions arise from this hypothesis. The first is that slower environmental change can restrict adaptation because all populations are finite and the best genotypes may be lost by genetic drift (Crow and Kimura 1970). The second prediction is that slower environmental change can limit adaptation by favoring the maintenance of similarly fit genotypes for longer periods, leading to a reduction in the mean population fitness and weaker selection for the genotypes with the highest fitness in the most extreme environments (Fisher 1930). Whether or not an adapting population has standing genetic diversity will profoundly affect the tempo and mode of evolution in changing environments (Matuszewski et al. 2015).

**Fig. 1.**
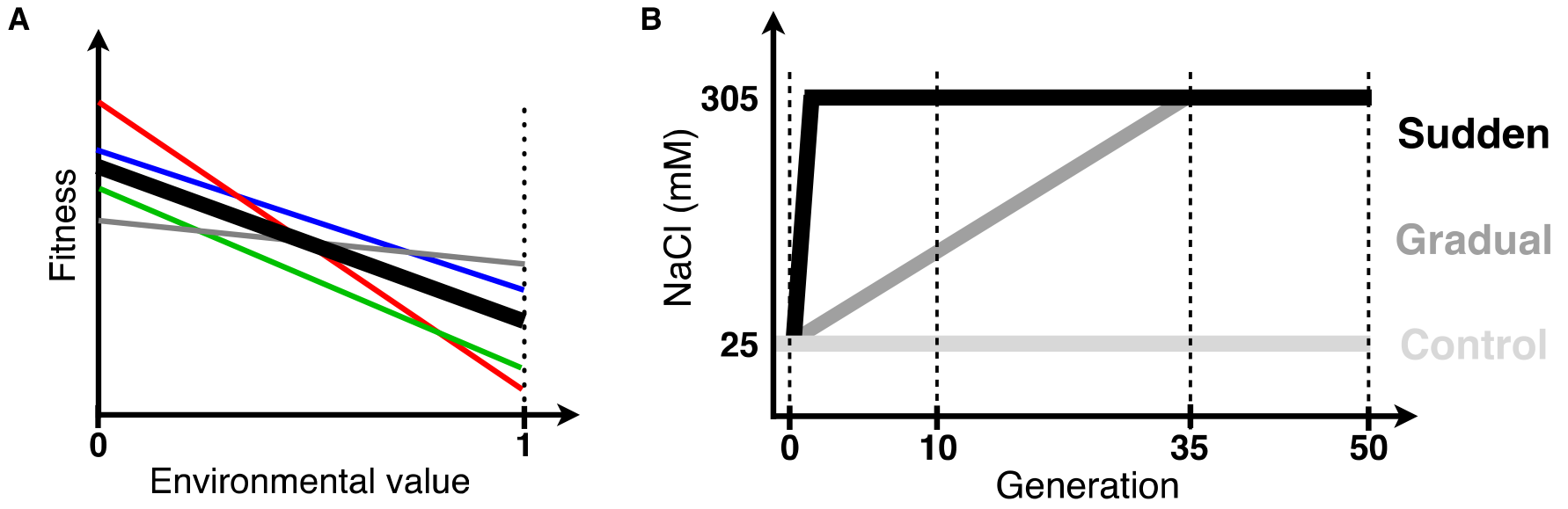
Fitness reaction norms and experimental evolution design. (A) Heritable genotype-by-environment fitness variance implies that genotypes (colored lines) have different growth rates along the value of the environmental factor(s) considered; known as “fitness reaction norms”, “tolerance functions” or “Finley & Wilkinson regressions” (Chevin et al. 2010, Walsh and Lynch 2014). Under density- and frequency-independent conditions, the relative difference of genotype growth rates to the average population growth rate (thick line) will determine the population genetic dynamics during evolution (Fisher 1930, Crow and Kimura 1970). Independently of their specific form, if there is crossing of fitness reaction norms particular genotypes will be favored at some environmental values while disfavored in others. For example, with a sudden change to an environmental value of 1 (vertical dotted line), from an ancestral environment 0, selection will favor the grey genotype, while a gradual change will initially favor the red genotype, then the blue one and only at a later period the grey genotype. (B) A 140-generation lab-adapted *C. elegans* population with genetic diversity, reproducing only by selfing, was the ancestor for experimental evolution. In the sudden regime, 4 replicate populations were faced from the first generation onwards to 305 mM NaCl in their growth media (black line). In the gradual regime, 7 replicate populations were faced with an 8 mM NaCl increase each generation until generation 35, being then kept at 305 mM until generation 50 (dark grey). In the control regime, 3 replicate populations were kept at 25 mM NaCl, the conditions to which the ancestor was adapted to (light grey). Vertical dashed lines indicate the time points where individuals were genotyped at single nucleotide polymorphisms (SNPs) across the genome (see also Fig. S1).

Here we show that heritable genotype-by-environment fitness variance is crucial for short-term adaptation under different rates of environmental change and illustrate the several population genetic mechanisms by which slower environmental change retards adaptation. To this end we performed experimental evolution at different rates of environmental change, using populations of the nematode *Caenorhabditis elegans* with standing genetic diversity where individuals can only reproduce by self-fertilization (Fig. 1B). At several time periods during experimental evolution we collected genome-wide single-nucleotide polymorphism (SNP) data at the individual level. We used these data, along with fitness data on the ancestral population, to develop a new inference model to understand the population genetics of adaptation to changing environments from standing genetic diversity.

## Results and Discussion

### Experimental evolution in changing environments

We performed replicated experimental evolution for 50 generations in the nematode *C. elegans* under different rates of change in the NaCl (salt) concentration that individuals experience from early larvae to adulthood (Fig. 1B and Table S1). In one regime, populations were suddenly placed in high salt concentration conditions (305 mM NaCl) while in another regime populations faced gradually increasing salt concentrations (see Materials and Methods). For the sudden regime, 4 replicate populations undergoing independent evolution were followed, while for the gradual regime we followed 7 replicate populations. All these populations are ultimately derived from a lab adapted ancestor population that has abundant genetic diversity (Chelo and Teotónio 2013, Noble et al. In Press), but where individuals reproduce exclusively by selfing and are expected to be homozygous at all loci throughout the genome (Crow and Kimura 1970, Theologidis et al. 2014). Except for salt concentrations, the same life-cycle of discrete and non-overlapping generations at stable census population sizes of 10^4^ individuals at the time of reproduction were maintained as during lab adaptation (see Materials and Methods). A control regime with 3 replicate populations was also maintained at the 25 mM NaCl conditions of lab adaptation. Given self-fertilization, the population sizes employed and the time span of experimental evolution neither mutation accumulation nor selection on new recombinants should contribute much to adaptation (Matuszewski et al. 2015, Teotónio et al. 2017).

### Modeling selection in changing environments

The fitness reaction norms are the key variables for predicting the evolutionary dynamics and the eventual outcome of adaptation in changing environments (Chevin et al. 2010) [Fig. 1A; see also chapter 44 in (Walsh and Lynch 2014)]. However, since fitness reaction norms are not directly available, one must resort to estimating them from the fitness and genotype data that we collected (see Materials and Methods; full inference model details in Supplementary Information). The data consist of individual-based biallelic SNP genotypes obtained at 3 time-points for each replicate population (Fig. 1B and Fig. S1), together with SNP genotypes and fitness data for the ancestral population. Our approach for modeling accounts for the genotyping setup, in which each individual was genotyped only in a pair of chromosomes (*C. elegans* is diploid with six chromosomes, for a genome size of 100 Mbp).

During experimental evolution, reproduction occurs exclusively by selfing, and so our model relies on effectively asexual population genetics dynamics. We consider deterministic environmental and population genetic dynamics, with discrete non-overlapping generations and viability selection. The environment faced in a given generation is represented by an environmental “value”, *x*, corresponding to the NaCl concentration. The population is composed of *G* selfing lineages, and we refer to the fitness reaction norm for a lineage *k* as *λ*_*k*_(*x*), corresponding to the expected number of live offspring produced under environment *x* (Fig. 1A). Each lineage is defined by a combination of haplotypes, based on the SNPs that were genotyped (Figs. S2 and S3). Inference is performed first by sampling the ancestral population (the lineages present and their starting frequencies), given the genotyping data, and then estimating the lineage reaction norms, repeating these two steps multiple times to obtain estimates of the reaction norm parameters. To estimate the lineage reaction norms, we assume they follow a specific parametric function of the environmental value, and then estimate the resulting parameters: here we consider log(*λ*_*k*_(*x*)) to be linear or quadratic functions of *x*.

### Experimental population genetics

Based on the genotyping data collected, we estimate that more than 200 distinct lineages are present in the lab adapted ancestral population (Fig. S2). The overwhelming majority of haplotypes observed are quickly selected against under all experimental evolution regimes (Fig. 2A and Fig. S4). We further find that populations faced with a sudden change in the first generation followed by constant high salt (305 mM NaCl) show for each region of the genome a single haplotype sweeping and nearing fixation by generation 50. In contrast, populations faced with a gradual increase in salt until generation 35 showed a different haplotype initially sweeping but then reverting in frequency when they were kept in the target high salt environment for another 15 generations.

**Fig. 2.**
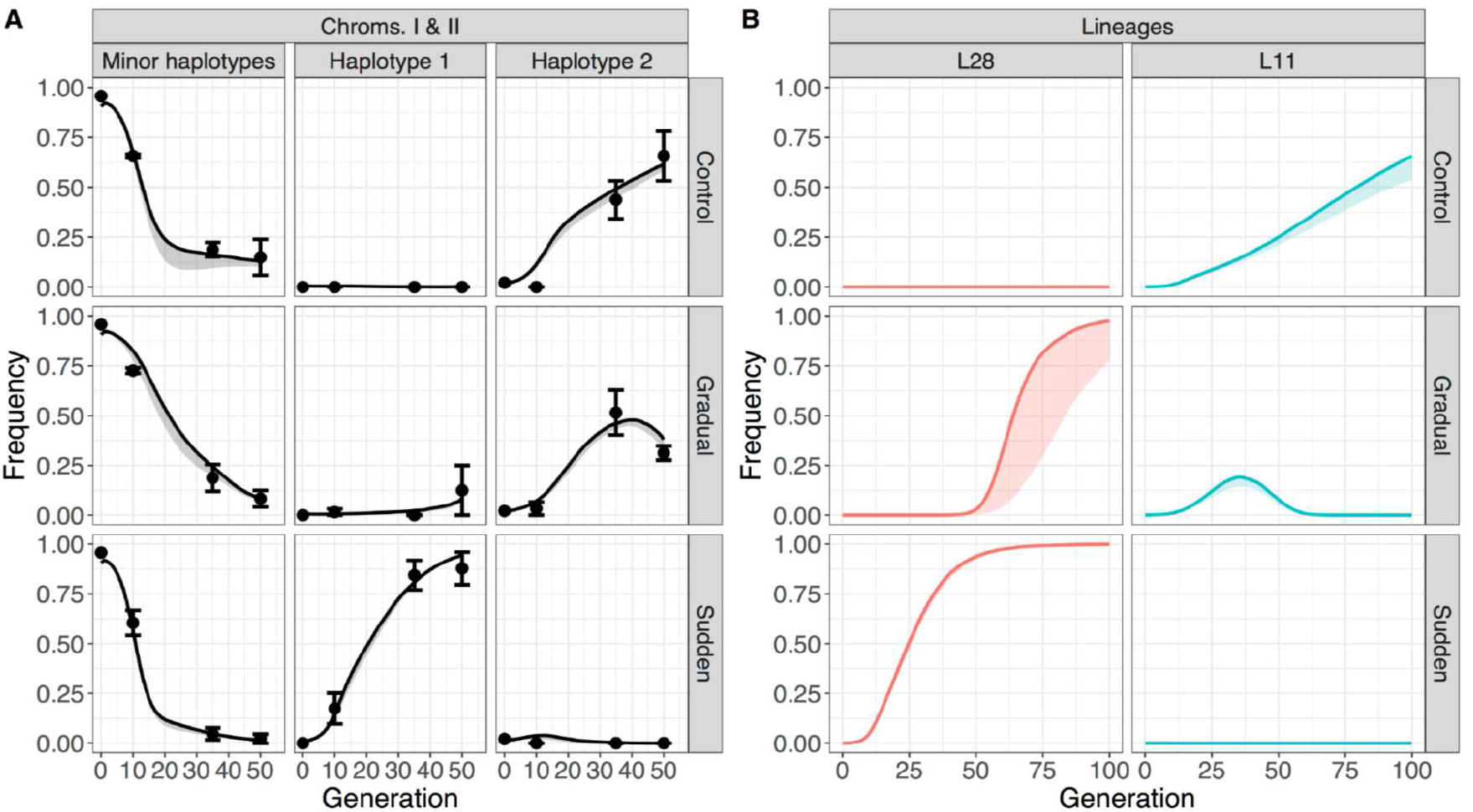
Observed and predicted experimental population genetic dynamics. (A) Left columns show that the majority of haplotypes observed for chromosomes I and II are quickly selected against in all experimental evolution regimes (rows). Middle and right columns show the two specific haplotypes in chromosomes I and II showing the greatest frequency change. Summary statistics and details of the pre-processing of the data for inference can be found in Figs. S2 and S3. Detailed results considering each replicate population are shown in Fig. S4. Symbols and error bars are the mean and one standard error of the observed haplotype frequencies among replicate populations. Line and shaded grey area are the trajectories inferred by modeling linear fitness reaction norms. (B) Inference results, evaluated over 100 generations (assuming that the gradual populations would be kept at high salt after generation 35), for the two main lineages (L28 and L11 in each column), assuming linear reaction norms. Detailed results for other lineages are shown in Fig. S5. Shaded colors correspond to confidence intervals considering the estimates obtained when sampling the ancestral population 20 times, with the line showing the median. Fig. S6 shows similar results when quadratic fitness reaction norms, instead of linear, are considered.

We initially modeled linear fitness reaction norms. The results indicate that the observed haplotype dynamics are consistent with a single lineage sweeping through the sudden populations (Figs. 2B, S4 and S5), which we name L28 (see below). Conversely, the gradual populations had an initial sweep of another lineage (L11), but then started to be overtaken by L28 by the end of the experiment. When modeling quadratic fitness reaction norms, the same conclusions are reached regarding haplotype and lineage dynamics (Fig. S6).

### Genotype-by-environment fitness interactions

Using whole-genome sequencing data on 100 lines derived from two gradual populations at generation 50 [from (Noble et al. In Press)], we identified the lines corresponding to the L28 and L11 lineages that we inferred (Fig S7 and Table S2). Our model predicts that the fitness reaction norms of these two lineages cross between 200-250 mM NaCl (Figs. 3A and S6C). To test this prediction, we directly assayed the fitness reaction norms of the ancestral population and the L28 and L11 lines. We find that the ancestral population fitness falls in-between those of the two lines at each salt level (Fig. 3B), and we find a close qualitative agreement with the model fit in that the line fitness reaction norms cross at about 225 mM (Fig. 3C). We also conducted head-to-head competitive fitness assays between L28 and L11, to account for any possible interactions that might not be apparent in the individual line growth assays. In these competition assays, performed for 2 generations, both lines were initially placed at 50:50 ratios. The results are remarkably similar to those under non-competitive conditions (Fig. 3D), indicating that interactions between the two lines are not significant.

**Fig. 3.**
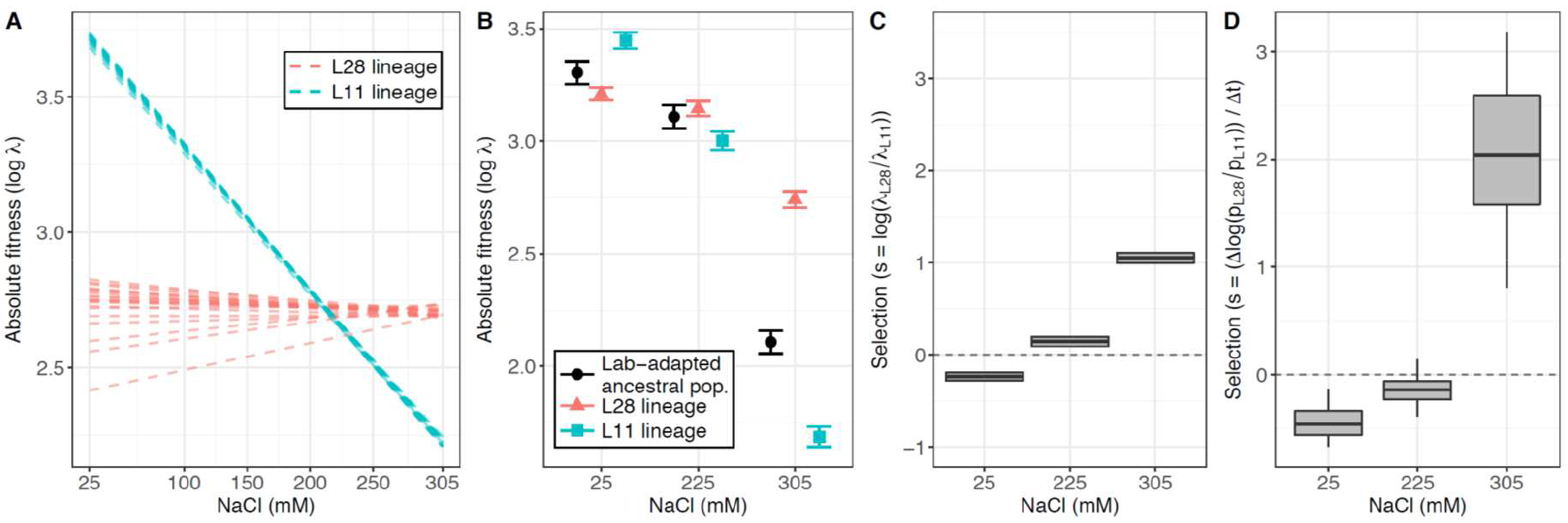
Crossing of fitness reaction norms. (A) Predicted fitness reaction norms of the L28 (red) and L11 (blue) lineages for 20 samples of the ancestral population (assuming linear reaction norms; see main text). (B) Absolute fitness of L28 and L11 lines, and the ancestor lab-adapted population at three salt concentrations (mean ± SE). (C) From (B), estimates (mean ± SE) of the expected relative fitness of L28 to L11 (selection coefficient) at three salt concentrations. (D) Similar to (C), but estimates from competitive fitness assays between L28 and L11. The two lineages were identified after genome-wide sequencing of 100 lines derived from two gradual populations at generation 50 (Fig. S7 and Table S2). Estimates were obtained using pooled-genotyping data on 18 SNPs that differ in L28 and L11 (see Materials and Methods, and Fig. S8 for calibration curves).

### Assessing how gradual environmental change affects adaptation to high salt

So far, our experiments and assays show that adaptation under different rates of environmental change is determined by the genotype-by-environment fitness variance present in the ancestral population. We next investigate the population genetic mechanisms by which this fitness variance can affect adaptation under slower environmental change.

We revived frozen stocks from the seven gradual populations at generation 35, the generation at which they reached the target high salt environment, and performed a new set of evolution experiments at two different population size regimes, 10^4^ and 2·10^3^, for 30 generations in constant high salt (Fig. 4A and Table S1). In this second set of experiments, we refer to each of the seven gradual populations as ancestrals. Two main factors, founder (bottleneck) effects and selection efficiency, could lead to differences in the evolutionary responses observed from each of the 7 new ancestral populations as well as between population size regimes. First, the best high salt lineage that we determined from the first set of evolution experiments, L28, was maintained at low frequencies during gradual evolution (Fig. 2B) and may have been lost before the second set of experiments started. The freezing and reviving process could also have resulted in L28 loss. If the L28 lineage was lost then future adaptation to high salt is compromised. The second factor is that the efficacy of natural selection on the best lineages may be lower because of increased genetic drift in small populations (Crow and Kimura 1970) or because of the maintenance of polymorphism in large populations (Fisher 1930), both mechanisms leading to lower selection efficiency.

**Fig. 4.**
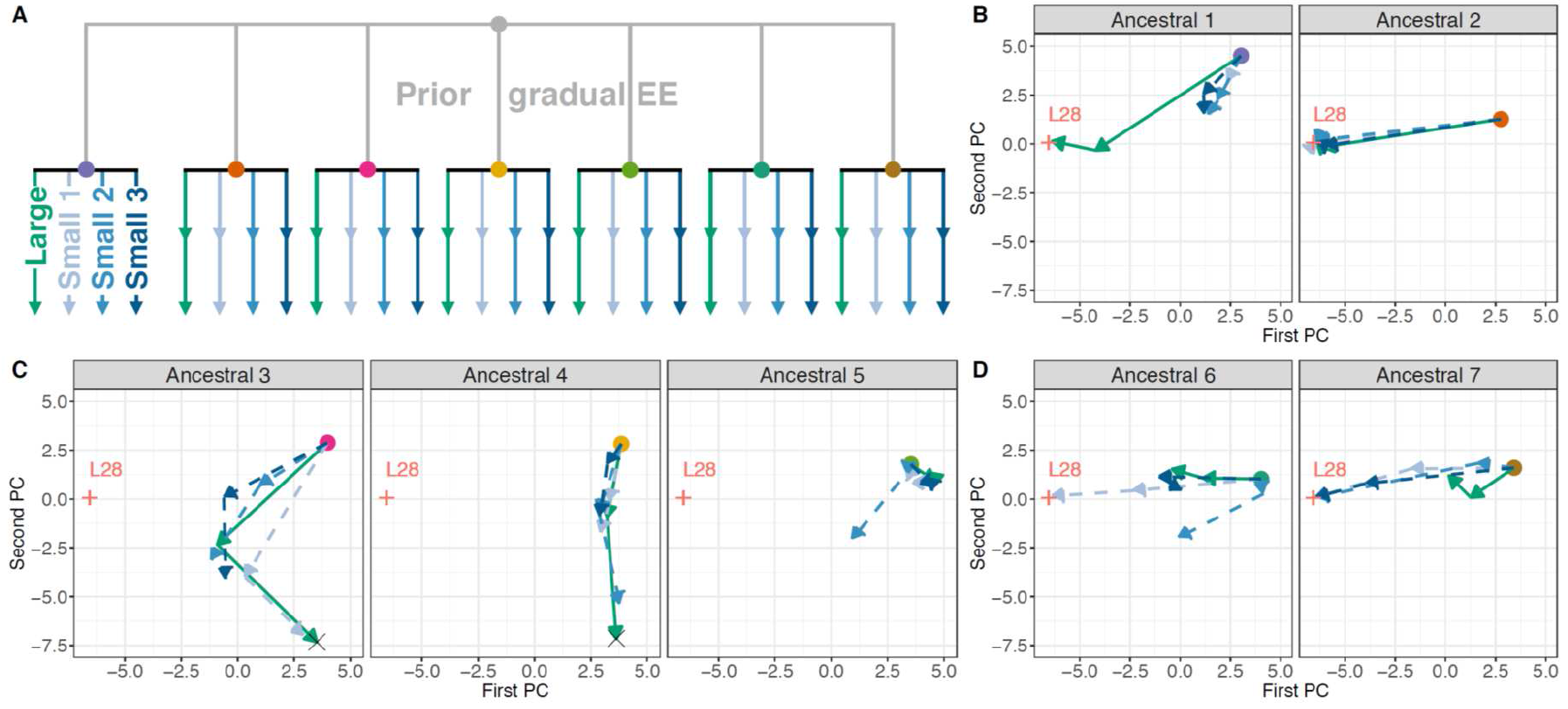
Gradual evolution restricts future adaptation to high salt. (A) Experimental evolution design at different population sizes. Seven gradual population at generation 35 become the ancestors (colored dots) for continued evolution in constant 305 mM NaCl for an extra 30 generations under large (green) and small (blue) population sizes. Populations were pool-genotyped after 15 and 30 generations. (B-D) Trajectories for the replicate populations under large and small population sizes, from the seven ancestor populations. These trajectories are based on principal component (PC) analysis of allele frequency data for 29 SNPs genotyped in pools of individuals, with the two first axis accounting for more than 70% of the variance (see also Fig. S9). Red crosses indicate the position of the L28 lineage, while the other crosses position another lineage. Analysis of the probability of a sweep by L28 is shown in Fig. S10.

### Genetic drift and selection efficiency

In two time points during this second set of evolution experiments, we genotyped a number of SNPs in pools of individuals, chosen to maximize the ability to distinguish lineage L28 (see Materials and Methods). We found that the evolutionary responses of the populations from the 7 ancestrals fell into three distinct categories.

The first category included two ancestral populations (Fig. 4B). From the first ancestral, it is clear that in large population sizes the L28 lineage swept and likely fixed, while at smaller population sizes the response was more constrained (Fig. 4B, and Figs. S8-S10). Despite population size differences, all populations derived from the second ancestral showed rapid sweeping of L28, indicating that L28 was initially at a high frequency. When comparing responses in this first category, we conclude that there was a founder effect, in that L28 was present at different initial frequencies, and that selection was more efficient when L28 was initially at high frequency and when evolution in high salt occurred at large population sizes.

In the second category, corresponding to three ancestrals, the L28 lineage was most likely lost before the second set of experiments (Fig. 4C). Strikingly, in ancestrals 3 and 4, another lineage, distinct from L28, swept more rapidly in large population sizes than in small population sizes, indicating again higher selection efficiency at larger population sizes. For the fifth ancestral, we can only conclude that the L28 lineage was lost before the second set of experiments.

Large population sizes are not, however, an assurance of higher adaptive rates in the high salt environment, as illustrated by the third category, consisting of two ancestrals (Fig. 4D). From them, we observed that the L28 lineage swept in a fraction of the populations, but exclusively in those with small population sizes. These results are consistent with a founder effect, in that initial evolution under the gradual regime maintained lineages that were either almost as fit as the L28 lineage (diminishing selection efficacy on L28), or that there was frequency-dependence between L28 and other lineages, e.g., (Chelo et al. 2013). Either way, some of the small populations must have lost these other competitive lineages (or maintained them at very low frequencies), before the second set of experiments, for the L28 lineage to sweep in them.

## Conclusions

The first set of experiments under different rates of environmental change demonstrates that adaptation depends on standing genotype-by-environment fitness variance (Figs. 1 and 2). This is a result that has been previously hinted in microbial evolution experiments that depended on mutation accumulation for adaptation (Lindsey et al. 2013, Gorter et al. 2015). Contrary to adaptation from pre-existing genetic diversity, however, when evolution occurs by the sequential fixation of mutations (and at the large population sizes usually employed in microbial evolution experiments), diminishing-returns epistasis for fitness appears to be involved for long-term adaptive dynamics to be predicted from short-term adaptive dynamics. In particular, short-term evolution must involve a sufficient number of generations so that adaptive gains become smaller with each new mutational event. Consistent with this scenario that diminishing-return epistasis for fitness makes evolution predictable, in constant environments, one yeast evolution experiment found a high degree of contingency in which specific mutations were sequentially fixed but not on how they interacted with each other at the fitness level (Kryazhimskiy et al. 2014).

Mutation accumulation experiments show that slower environmental changes allows more time for the exploration of mutational space and the possibility to fix mutations at intermediate environments that predispose subsequent fixation of additional epistatic mutations at more extreme environments. In (Gorter et al. 2015), under some stressors, slow environmental change retarded adaptation but not the fitness gains in the most extreme environments, and in (Lindsey et al. 2013) the populations that survived a sudden environmental change had higher fitness than those that survived a more gradual change, suggesting, just as in our experiments from standing genetic diversity, a key role of genotype-by-environment fitness interactions. Some authors refer to this phenomenon as “environmental epistasis” since the non-additive fitness interactions between fixed mutations are themselves environmentally-dependent (Remold and Lenski 2004). With standing genetic variation, we showed that genotype-by-environment fitness interactions are sufficient to explain adaptive dynamics, independently of non-additive interactions between competing genotypes, such as, in our case, negative frequency-dependence (Fig. 3). The emerging picture is that genotype-by-environment fitness interactions are critical for adaptation to changing environments when evolution occurs from standing genetic diversity, and that both genotype-by-environment interactions and epistasis are important when evolution occurs from mutation.

Little theoretical work has focused on understanding the population genetics of adaptation from pre-existing genetic diversity. An exception is the study by Matuszewski and colleagues (Matuszewski et al. 2015), which explored the distribution of fitness effects of fixed alleles starting from standing variation under a moving trait under stabilizing selection. They found that populations facing a fast environmental change show larger trait changes than those facing a slow environmental change, due to increases in both the expected number of fixations and the expected trait effect per allele substitution. Although they did not analyze situations of complete linkage, as in our evolution experiments, they nonetheless predicted a higher number of fixations under faster environmental change, and that adaptation would be deterred under slower environmental changes. We found with the continued evolution experiments in high salt that slower environmental change will indeed maintain polymorphism (Fig. 2) and compromise adaptation (Fig. 4). This is because small population sizes and bottlenecks reduce the efficacy of selection on the best genotypes, and/or promote their loss, and the maintenance of polymorphism for long periods in large populations reduces the likelihood of the single best genotype becoming fixed. Besides being remarkably consistent with the predictions of (Matuszewski et al. 2015), these findings confirm classical theory on the role of genetic drift in the loss of adaptive diversity and on the role of standing genetic variation in reducing mean population fitness (Fisher 1930, Crow and Kimura 1970). They also indicate that long-term adaptation (say, for 100 generations, as in our model in Fig. 2B) cannot be readily predicted from short-term adaptation (say, from the first 35 generations, as in our first set of experiments). Except in the most contrived cases where the identity and relative frequency of ancestral genotypes is known *a priori* because of being constructed, e.g., (Gresham et al. 2011), long-term adaptation to changing environments in natural populations is not likely to be well predicted from short-term adaptation; but see (Charmantier et al. 2008).

From an empirical perspective, the inference model that we developed here, where only partial information about the short-term evolutionary trajectories of fitness and genetic diversity is used, is a significant step in understanding evolution in changing environments. Using our approach, for natural populations, partial genomic and fitness observations may allow predicting adaptation to changing environments and, possibly, the likelihood of extinction. Although most natural species are sexual and thus recombination of pre-existing diversity could play a role in adaptation to changing environments, it is unlikely that new recombinants will contribute much given the limited population sizes of most species, and the short time spans of environmental change relative to their generation times, but see, e.g., (Aggarwal et al. 2015). Related modeling approaches to ours have been proposed to predict, for example, the within-host evolution of influenza from single infection events (Illingworth et al. 2014). In our case, we add the possibility for environmental change. In the agricultural literature similar approaches have also been devised to predict plant yield and animal production in several environments, but either the environment is modeled as discrete or linear fitness reaction norms are usually considered, and it is still unclear how to best model the heritability of fitness from genomic data, cf. (Gomulkiewicz and Kirkpatrick 1992, Walsh and Lynch 2014). Including stochasticity in the population genetics and recombination would improve out method and allow explicit predictions of the loss of genetic variance in the fitness reaction norms under gradual environmental change.

## Materials and Methods

Detailed materials and methods can be found in the Supplementary Information file.

### Experimental evolution in changing environments

All populations employed are ultimately derived from a hybrid population of 16 wild isolates (Teotónio et al. 2012), followed by 140 generations of laboratory domestication to a 4-day non-overlapping life-cycle under partial self-fertilization (selfing) at census sizes of N = 10^4^ (Teotónio et al. 2012, Chelo et al. 2013), and finally introgression and homozygosity of the *xol-1 (tm3055)* sex determination mutant allele at high populations sizes for 16 generations to generate an ancestral population only capable of reproduction by selfing (Theologidis et al. 2014). For experimental evolution in changing environments (Fig. 1B), ancestral population samples were thawed, expanded in numbers and first larval staged (L1s) seeded at the appropriate densities to three regimes. The sudden regime was characterized by the same conditions to which previous lab-adaptation occurred, except that the NGM-lite media (US Biological) where worms grew was supplemented with NaCl (305 mM) from the start and for 50 generations (4 replicate populations; Supplementary Information, Table S1). For the gradual regime plates were supplemented with increasing concentrations of NaCl from 33 mM at generation 1 to 305 mM NaCl at generation 35 and onwards until generation 50 (7 replicate populations). A control regime was maintained in the ancestral environmental conditions without any salt supplement (3 replicates).

### Experimental evolution at different population sizes

All 7 replicate populations from the gradual regime at generation 35 were revived from frozen stocks, expanded in numbers for two generations, and then split into two regimes: large population sizes of N=10^4^ and small population sizes of 2·10^3^. From each of the seven gradual populations at generation 35, one replicate was maintained at large population sizes and three replicates were maintained at small population sizes. All populations were kept at constant 305 mM NaCl for 30 generations. Over 10^3^ L1s were collected per population at generation 15 and 30, for pool-genotyping.

### Fitness assays

The ancestral population (before salt adaptation) was thawed from frozen stocks and individuals reared for two generations at 25 mM before they were exposed to the three assay NaCl treatments (Fig. 3B). On the third generation, five Petri dishes per NaCl treatment were seeded with 10^3^ L1s per plate. These five plates constituted one technical replicate, and there were four of these for each salt treatment. After 66 h, individuals were harvested and exposed to a 1 M KOH:5% NaOCl solution (to which only embryos survive). After 16 h, debris was removed and the total number of live L1s in each tube was estimated by scoring the number of L1s. For analysis, the per-capita L1-to-L1 growth rate values were linearly modelled in R (R Development Core Team 2015): `log (growth_rate) ~ salt_treatment`. Least-square estimates were obtained using the R package *lsmeans* (Lenth 2015).

During experimental evolution in changing environments, one lineage swept through the sudden populations, while another lineage was initially sweeping though the gradual populations when they were at intermediate salt concentrations (Fig. 2B). From two gradual populations at generation 50, we derived in (Noble et al. In Press) 100 lines which were whole-genome sequenced. Of these, we identified lines L28 and L11 as representatives of the lineages predicted to explain the experimental population dynamics. For them, fitness assays were conducted as for the ancestral population, for two full generations (Fig. 3BC), over three blocks (defined by when L28 and L11 were revived from frozen stocks). For analysis, we used a mixed effects model (Bates et al. 2015) via the R formula: `log(growth_rate)~salt_treatment * line + assay_generation*line + (1|block)`. To estimate the expected selection coefficient of L28 relative to L11 we used *lsmeans* formulation: `pairwise ~ line | salt_treatment`.

L28 and L11 were also assayed in head-to-head competitions (Fig. 3C). They were thawed from frozen stocks and reared for two generations at 25 mM NaCl before they were set up at three NaCl concentrations: 25 mM, 225 mM and 305 mM. On the third generation, L1 larvae from the two lineages were mixed in 1:1 ratio, at a density of 10^3^ L1s in each of two Petri dishes per replicate assay. Each replicate assay was maintained for two generations. At both the assay generations, L1 samples were collected for pool-genotyping of single nucleotide polymorphisms (SNPs). Assays were performed in three blocks, with 3 replicate populations per salt concentration in each of two blocks, and 4 replicate populations in the third block. The data for analysis was based on the L28 and L11 SNP frequency values obtained after doing calibration curves where the ratio of both lines was known. For analysis, the estimated frequencies for L28 were forced to be in the interval (0.005, 0.995). To estimate the relative selection coefficients we again used a mixed effects model: `log(Odds_Ratio_L28) ~ salt_treatment * assay_generation + (1|Techn_replicate)`.

### Genotyping

Individual L4 genomic DNA was prepared with the ZyGEM prepGEM TM Insect kit following (Chelo and Teotónio 2013). A total of 925 biallelic SNPs across the genome were assayed by iPlex Sequenom™ MALDI-TOF methods (Bradic et al. 2011). We chose the SNPs that we knew were segregating in the lab-adapted population (Noble et al. In Press). Due to the limited amount of genomic DNA, each individual was assayed for two of the six *C. elegans* chromosomes, each pair of chromosomes being referred to as a region (chromosomes I and II: region 1; III and IV: region 2; V and VI: region 3). For genotyping, larvae at the L4 (immature) stage: 64 larvae per region, from the ancestral M00; from each of the evolved populations (generations 10, 35 and 50), 16 L4s were sampled per region. Briefly, quality control was based on discarding SNPs with a high frequency of heterozygous calls, SNPs with a high frequency of genotyping failures (> 30%), and individuals in which many SNPs failed genotyping (> 25%). The 761 SNPs that passed quality control were imputed into chromosome-wide haplotypes using fastPHASE (Scheet and Stephens 2006). These SNPs were evenly spaced at an average of 0.30-0.38 cM, according to the genetic distance of (Rockman and Kruglyak 2009). Number of individuals per population, after quality control, can be found in Fig. S1B.

Genomic DNA from pooled samples was prepared using the Qiagen Blood and Tissue kit, and genotyped for 84 SNPs in chromosomes I, IV and V, using the iPlex Sequenom methods in 3 technical replicates for each SNP assay. In parallel, pooled gDNA was prepared to calibrate SNP L28 allele frequencies when mixed with L11 or the ancestor population at several known proportions (8-14 technical replicates each). After quality control, we retained 29 SNPs, 18 of which differentiating L28 and L11 (Fig. 3D). We interpolated expected L28 frequencies from the calibration curves, using Levenberg-Marquardt algorithm in R package *minpack.lm* (Elzhov et al. 2016). For the principal component analysis of the matrix containing the frequency of the alternative alleles in each sample (Fig. 4), the function prcomp in R was used.

### Fitness reaction norms

We assume an effectively asexual population genetics model for a haploid organism, ignoring segregation within loci and recombination among loci. The model also considers deterministic environmental and population dynamics, discrete non-overlapping generations and viability selection, with the only environmentally-relevant variable being the NaCl concentration. We assume an infinite population size, such that any given lineage never goes extinct (although the frequency may become very small), that there are no density- or frequency-dependencies, and that trans-generational effects are absent.

A population is composed of *G* lineages, such that the frequency of the *k*-th lineage in generation *t* + 1, denoted by 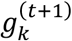, is given by:

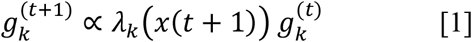

where *x*(*t*) is the environment value faced in generation *t*, and *λ*_*k*_(*x*) the expected number of live offspring produced by lineage *k* when faced with the environment *x*. In this way, the function *λ*_*k*_(*x*) corresponds to the fitness reaction norm for lineage *k*.

Following the setup used for genotyping, the genome is divided into *L* non-overlapping regions, and we refer to the haplotype in a region as a region-wide haplotype (RWH). A “lineage” *k* is described by a tuple *S*_*k*_, indicating the RWHs in each region, such that *S*_*k*_ = (*l*_*k*,1_, *l*_*k*,2_, ···, *l*_*k*,*L*_). We assume that the fitness reaction norm of a lineage is an additive function of the fitness reaction norm of the RWHs in that lineage:

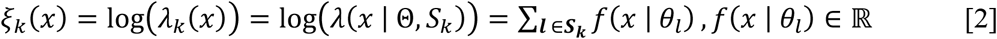

where Θ is a vector of parameters for all the region-wide haplotypes, *θ*_*l*_ the parameters for RWH *l*, and *f*(*x* | *θ*_*l*_) the parametric function describing the fitness reaction norm for a single RWH. We considered *f*(*x* | *θ*_*l*_) to be a linear (*f*(*x* | *θ*_*l*_) = *a*_*l*_ *x* + *b*_*l*_, such that *θ*_*l*_ = (*a*_*l*_, *b*_*l*_)) or quadratic function (*f*(*x* | *θ*_*l*_) = *a*_*l*_ *x*^2^ + *b*_*l*_ *x* + *c*_*l*_, such that *θ*_*l*_ = (*a*_*l*_, *b*_*l*_, *c*_*l*_)) of the environmental value *x*.

Given genotyping and/or fitness data at *H* time-points plus the ancestral, we consider distinct epochs of the experimental evolution, evaluated at generations *T*_0_, *T*_1_, ···, *T*_*H*_ (such that *T*_0_ = 0, *T*_1_ = 10, *T*_2_ = 35 and *T*_3_ = 50; Fig. 1B). To denote the epoch to which a certain variable corresponds, a superscript inside square brackets is used. For a single population, the frequency of lineage *k* in epoch *ℎ*, denoted by 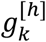, follows from the frequencies of the lineages in the previous epochs:

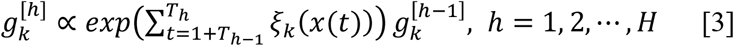

where *x*(*t*) is the environment faced in generation *t*. The ancestral population, consisting of *G* lineages, is described by two variables: *A* = (*S*_1_, *S*_2_, ···, *S*_*G*_), corresponding to the RWHs
present in each lineage; and 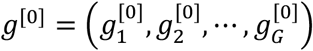, specifying the frequency of each lineage (such that 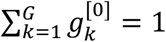).

### Inference

For inferring the lineage fitness reaction norms, *λ*_*k*_(*x*), we consider that *A* and *g*^[0]^ are known. Since this is not the case in the analysis of the experimental data, we sample the pair (*A*, *g*^[0]^), given the experimental data, and then estimate the RWH parameters Θ, repeating these two steps multiple times (sections 1.7.6 and 1.7.7 of the Supporting Information).

Under the population genetics model used, all replicate populations within a single evolutionary regime *c* have the same dynamics of the lineage frequencies 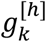. Let 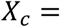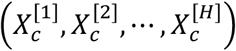 denote the sequence of environmental values in regime *c*, where 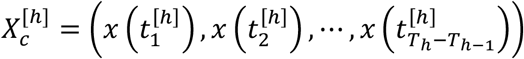, 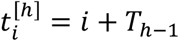. Inference is framed in a maximum likelihood context, with contributions from each evolutionary regime, given the fitness and genotyping data. We consider without loss of generality that fitness and genotyping data are available for all epochs *T*_0_, *T*_1_, ···, *T*_*H*_ for each regime. The case in which data is available only for certain epochs is treated by evaluating the corresponding likelihood function only for those epochs. The Supporting Information details how the input data, at the level of the replicate populations, is converted to that at the level of each regime.

Let 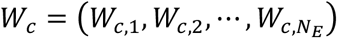 denote the fitness data on regime *c*, with *N*_*E*_ assay environments, with *x*_*m*_ being the environmental value, and 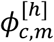 the observed population-averaged fitness value of a population from regime *c* in epoch *ℎ* in the *m*-th assay environment. We assume a log-normal model for noise in the observed values 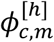. The log-likelihood for the RWH parameter vector Θ given the fitness data on regime *c* is then:

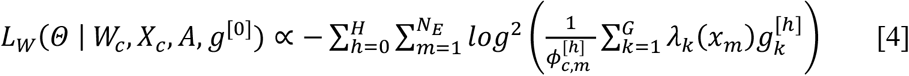

Let 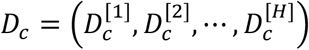 be the genotyping data on regime (note that it does not include the data on the ancestral), such that 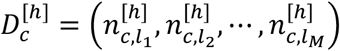, where 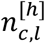 is the number of copies of RWH *l* that were observed in epoch *ℎ* in regime *c*. Then, the log-likelihood given the genotyping data on regime c is given by:

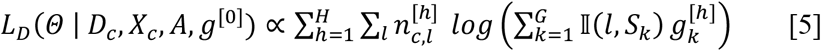

where 𝕀(*l*, *S*_*k*_) is an indicator function, equal to 1 if lineage *k* has RWH *l*, or equal to 0 otherwise.

Considering all evolutionary regimes *C*, the log-likelihood is then obtained by combining equations [4] and [5]:

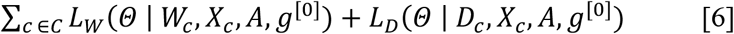

Model fitting is then performed by maximizing equation [6], using a gradient-based optimization algorithm, starting from random initial conditions.

## Acknowledgments

We thank J. Costa, A. Crist, H. Gendrot and I. Theologidis for support with nematode handling and sample preparation, L. Noble for help with the genomic data analysis, and the Center for Scientific Computing from the CNSI, MRL, at UC Santa Barbara, an NSF MRSEC (DMR-1121053) and NSF CNS-0960316 supported facility, for computation. We thank R. Gomulkiewicz, J. Hermisson, M.-A. Félix, S. Matuszewski and L. Noble for discussion. S.D. is a fellow of the Labex MemoLife (ANR-10-LBX-54 MEMO LIFE and ANR-IDEX-0001-02-PSL). Financial support from the National Science Foundation (EF-1137835) to S.R.P., the Human Frontiers Science Program (RGP0045/2010), the European Research Council (FP7/2007-2013/243285) and Agence Nationale de la Recherche (ANR-14-ACHN-0032-01) to H.T. All data and code for analysis will be deposited in public repositories.

